# Functional evolution of a bark beetle odorant receptor clade detecting monoterpenoids of different ecological origins

**DOI:** 10.1101/2020.12.28.424525

**Authors:** Xiao-Qing Hou, Jothi Kumar Yuvaraj, Rebecca E. Roberts, Dan-Dan Zhang, C. Rikard Unelius, Christer Löfstedt, Martin N. Andersson

## Abstract

Insects detect odors using an array of odorant receptors (ORs), which may expand through gene duplication. How specificities evolve and new functions arise in related ORs within a species remain poorly investigated. We addressed this question by functionally characterizing ORs from the Eurasian spruce bark beetle *Ips typographus*, in which antennal detection and behavioral responses to pheromones, volatiles from host and non-host trees, and fungal symbionts are well described. In contrast, knowledge of OR function is restricted to two receptors detecting the pheromone compounds (*S*)*-*(–)-ipsenol (ItypOR46) and (*R*)-(–)-ipsdienol (ItypOR49). These receptors belong to a species-specific OR-lineage comprising seven ItypORs. To gain insight into the functional evolution of related ORs, we characterized the five remaining ORs in this clade, using *Xenopus* oocytes. Two receptors responded primarily to the host tree monoterpenes (+)-3-carene (ItypOR25) and *p*-cymene (ItypOR27). Two receptors responded to oxygenated monoterpenoids produced in relatively large amounts by the beetle-associated fungi, with ItypOR23 specific for (+)-*trans*-(1*R*,4*S*)-4-thujanol, and ItypOR29 responding to (+)-isopinocamphone and similar ketones. ItypOR28 responded to the pheromone *E*-myrcenol from the competitor *Ips duplicatus*. Overall, the OR responses match well with those of previously characterized olfactory sensory neuron classes except that neurons detecting *E-*myrcenol have not been identified. The characterized ORs are under strong purifying selection and demonstrate a shared functional property in that they all primarily respond to monoterpenoids. The variation in functional groups among OR ligands and their diverse ecological origins suggest that neofunctionalization has occurred early in the evolution of this OR-lineage following gene duplication.

## Introduction

Olfaction is of utmost importance to the ecology of insects, with odors mediating fitness-related activities such as mate choice and the search for food and oviposition sites, and maintenance of symbioses with microbes (Hansson and Stensmyr, 2011; Andersson et al., 2015; Fleischer et al., 2018; Kandasamy et al., 2019). Odors are detected by an expansive gene family encoding odorant receptors (ORs), which are expressed in the olfactory sensory neurons (OSNs) in the insect antennae (Clyne et al., 1999). As such, the ORs are crucial to insect life and they underlie the diverse chemical ecologies, adaptations, niche specializations, and evolutionary divergence seen in the Insecta class, which dominates most habitats of our planet. Yet, the functional evolution of this gene family is poorly understood (but see e.g., Ramdya and Benton, 2010; de Fouchier et al., 2017; Yuvaraj et al., 2017; Guo et al., 2020), and little is known with regards to how new functions evolve in these receptors, especially in non-model insects. This applies to both the molecular mechanics that underlie specificity changes (but see Leary et al., 2012; Hopf et al., 2015) and the ecological selection pressures that drive the evolution of the ORs.

New OR genes originate through gene duplication. Hence, the duplicated receptors that are retained and expressed will initially have the same odor specificity as their parental ORs, but may later acquire neutral or adaptive mutations that alter their specificity (Nei et al., 2008; Andersson et al., 2015). One may therefore predict that duplicated “offspring” OR paralogues are likely to adopt similar functions as their “parent” ORs, i.e., detecting compounds with shared chemical characteristics (but see also Adipietro et al., 2012). In this study, we investigated the functional evolution of OR paralogues within a species-specific OR-radiation from the Eurasian spruce bark beetle *Ips typographus* L. (Coleoptera; Curculionidae; Scolytinae).

As keystone species in forest ecosystems, bark beetles play an important role in the decomposition of wood and recycling of nutrients (Edmonds and Eglitis, 1989; Raffa et al., 2016). However, a minority of bark beetle species are able to kill healthy trees during outbreak periods, which has become an increasing threat to conifer forests, causing great economic loss (Raffa, 2001; Kausrud et al., 2011; Raffa et al., 2016; Biedermann et al., 2019). In Eurasia, *I. typographus* is considered the most serious pest of Norway spruce (*Picea abies*) (Økland and Bjørnstad, 2003). When *I. typographus* population density surpasses a critical threshold, the defense of healthy trees can be overcome through mass-attacks, and entire forest landscapes can be quickly transformed (Wyatt, 2014; Raffa et al., 2016). The mass-attacks are coordinated by a male-produced aggregation pheromone consisting of (4*S*)-*cis*-verbenol and 2-methyl-3-buten-2-ol, attracting both sexes (Schlyter et al., 1987). The attraction is, however, inhibited by several other pheromone compounds produced by *I. typographus* during the later attack phases or by heterospecific beetles, including *E-*myrcenol and specific enantiomers of verbenone, ipsenol, and ipsdienol (Francke et al., 1980: Birgersson et al., 1984; Schlyter et al., 1989; 1992).

In addition to the beetle-produced semiochemicals, the pheromone attraction is modulated by volatiles from non-host trees and defense compounds from the host (Zhang and Schlyter, 2004; Andersson et al., 2010; Schiebe et al., 2012; Binyameen et al., 2014; Unelius et al., 2014). Recent laboratory studies have also demonstrated the importance of volatiles in the maintenance of symbioses between *I. typographus* and its fungal associates, with beetle preferences for certain fungi being mediated via olfactory cues (Kandasamy et al., 2019). Once inoculated in a tree, these fungi may provide nutrients to maturing beetles by direct feeding (Kandasamy et al., 2019), metabolize host tree defenses (Kandasamy, 2019), and possibly accelerate tree death (Horntvedt et al., 1983). Extensive efforts have been made to understand the ecological roles of *I. typographus*-associated compounds and to characterize the OSNs that specifically detect them. To date, a total of 23 OSN classes have been reported. The key ligands of these neurons include pheromone compounds from conspecific and heterospecific bark beetles, volatiles from host and non-host plants, and compounds produced by fungal symbionts (Mustaparta et al., 1984; Tømmerås et al., 1984; Tømmerås, 1985; Andersson et al., 2009; Andersson 2012; Raffa et al., 2016; Kandasamy, 2019; Kandasamy et al., 2019; Schiebe et al., 2019; Zhao et al., 2019).

The odor selectivity of an OSN depends on the characteristics of the OR that is expressed in that neuron. Hence, functional characterization of ORs is important for understanding the evolution of olfactory specialization. Despite being the largest insect order, functional information of ORs from Coleoptera is, however, limited compared with moths (e.g., Grosse-Wilde et al., 2007; Zhang and Löfstedt, 2015; Zhang et al., 2016; de Fouchier et al., 2017; Guo et al., 2021), flies (e.g., Hallem and Carlson, 2006; Mansourian and Stensmyr, 2015) and mosquitos (Carey et al., 2010; Wang et al., 2010). To our knowledge, only seven beetle ORs have been characterized so far, of which two belong to *I. typographus* (Mitchell et al., 2012; Antony et al., 2020: Mitchell and Andersson 2020; Yuvaraj et al., 2021; Wang et al., 2020). In our previous study, the two *I. typographus* ORs (ItypOR46 and ItypOR49), responded specifically to the pheromone compounds (*S*)-(−)-ipsenol and (*R*)-(−)-ipsdienol respectively. These ORs are part of an *Ips-*specific OR-lineage radiation comprising seven ORs (Yuvaraj et al., 2021). This radiation is present within the coleopteran OR subfamily named Group 7, which is highly expanded in the Curculionidae family (Andersson et al., 2013; Andersson et al., 2019; Mitchell et al., 2020). The other five ORs (ItypOR23, OR25, OR27, OR28 and OR29) in this clade did not respond to any stimulus when tested in HEK293 cells, which was likely due to insufficient protein levels (Yuvaraj et al., 2021). Because this clade represents the largest highly-supported OR-radiation in the antennal transcriptome of this species and many of the OR genes are highly expressed (Yuvaraj et al., 2021), we hypothesize that these ORs have key ecological functions. Thus, targeting the five remaining ORs using a different expression system could provide information on the functional evolution of ORs in this species and also shed light on the general question of how odor specificities may evolve among OR paralogues within species-specific radiations and how their functions relate to species ecology. Hence, the primary aim of this study was to functionally characterize these five ORs using *Xenopus* oocytes and a large panel of ecologically relevant compounds. We asked whether the clade houses additional ORs specifically detecting other bark beetle pheromone compounds, or whether the ORs detect structurally similar chemicals (i.e., oxygenated monoterpenoids) of different biological origins and ecological meanings. The second aim was to link the responses of the functionally characterized ItypORs with the responses of previously identified OSN classes.

## Results

### *Characteristics of the seven ORs in the* Ips*-specific clade*

The five ItypORs targeted for functional characterization are part of an *Ips*-specific OR-lineage comprising seven ItypORs, and this lineage belongs to the monophyletic OR Group 7 in beetles (Mitchell et al., 2020; Yuvaraj et al., 2021). The phylogenetic relationships of these ORs and a subset of additional Group 7 ORs from *I. typographus* and the mountain pine beetle *Dendroctonus ponderosae* (‘Dpon’) are shown in Fig. 1A. The expression levels (relative to Orco; data from Yuvaraj et al., 2021) and proposed key ligands of the seven ItypORs (based on our oocyte recordings; see below) are shown in Fig. 1B, D. Except for ItypOR27, these OR genes are all among the top 20 (out of 73) most highly expressed OR genes in the antennae of this species (Yuvaraj et al., 2021). The pair-wise amino acid identity amongst the seven ORs ranges from 36.6 % to 49.9 % (Fig. 1C).

**Fig. 1.**
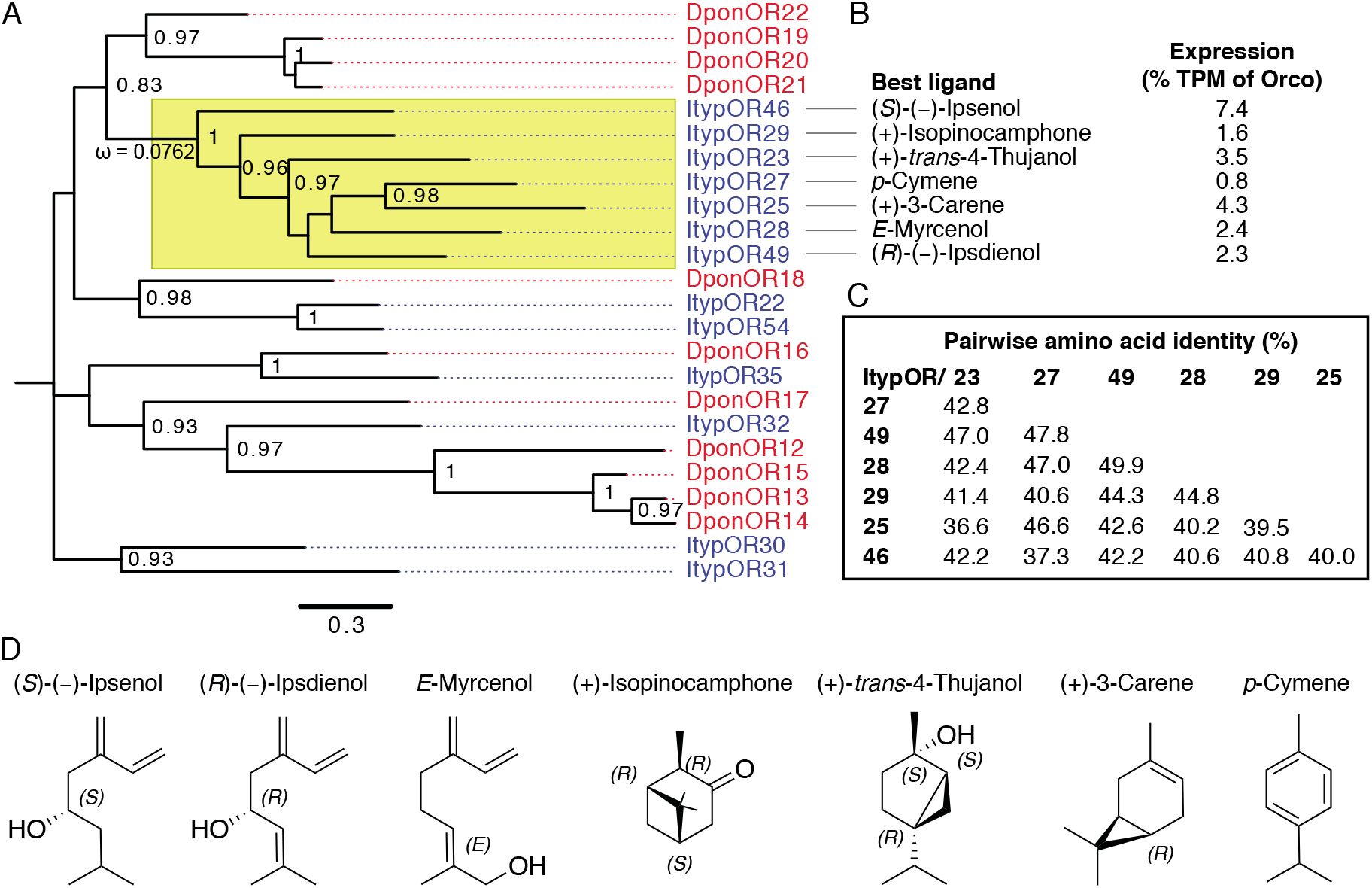
(A) Unrooted maximum likelihood phylogenetic tree showing the relationships among select Group 7 (Mitchell et al., 2020) odorant receptors (ORs) from *Ips typographus* (‘Ityp’; blue) and *Dendroctonus ponderosae* (‘Dpon’; red). The tree is based on a MAFFT alignment of amino acid sequences and constructed using FastTree 2.1.11. The clade containing the seven ItypORs is highlighted in yellow and the strong purifying selection (ω = 0.0762) is indicated on the branch. Numbers at nodes are local support values, calculated using the Shimodaira-Hasegawa (SH) test implemented in FastTree (SH values below 0.7 are not shown). (B) The primary ligands and expression levels (transcripts per million; TPM) relative to Orco of the seven *Ips*-specific ORs (data from Yuvaraj et al., 2021). (C)The pair-wise amino acid identity amongst the seven ORs, based on a MAFFT alignment. (D) Chemical structures of the main ligand for each of the seven ItypORs in the clade.

We evaluated the selective pressure acting on the seven ItypORs by calculating the ratio of nonsynonymous to synonymous substitutions (dN/dS or ω) using PAML (Yang, 1997; Yang and Nielsen, 1998). We first tested whether the clade has a uniform dN/dS ratio (one ratio model, M0) or variable ratios on different branches (free ratio model, M1) by likelihood ratio test (LRT). The result showed that the one ratio model could not be rejected (*P* = 0.19), and the dN/dS is 0.07623 (<< 1) for this clade. We next examined whether there are positively selected sites across the sequences using an LRT, comparing the NSsites models M7 (beta) and M8 (beta and ω). The results showed that the M7 model could not be rejected (*P* = 0.93), and no positively selected sites were reported across the sequences. Taken together, our dN/dS analysis indicate that the seven ItypORs are under strong purifying selection.

### ItypOR23 and ItypOR29 respond to volatiles mainly produced by fungal symbionts

Screening experiments with *Xenopus* oocytes co-expressing ItypOR23 and ItypOrco showed a primary response to (+)-*trans*-4-thujanol, and secondary responses to (±)-3-octanol, (–)-terpinen-4-ol, as well as acetophenone (Fig. 2A and B). Dose-response experiments showed that ItypOR23 is more sensitive to (+)-*trans*-4-thujanol than to the other three compounds (Fig. 2C), with this compound eliciting larger responses at each tested concentration. Two recent studies (Schiebe et al., 2019; Kandasamy, 2019) using partly overlapping odor panels characterized an OSN class specific for (+)-*trans*-4-thujanol (named OSN class tMTol), with responses very similar to those of ItypOR23 (Table 1). Although a weak secondary response to 1-octen-3-ol was observed in the OSN (Kandasamy, 2019) but not in the OR, and a weak secondary response to acetophenone in the OR but not in the OSN, the high similarity in the responses to the three best ligands suggests that ItypOR23 is likely to be the matching OR for this OSN class.

**Table 1.**
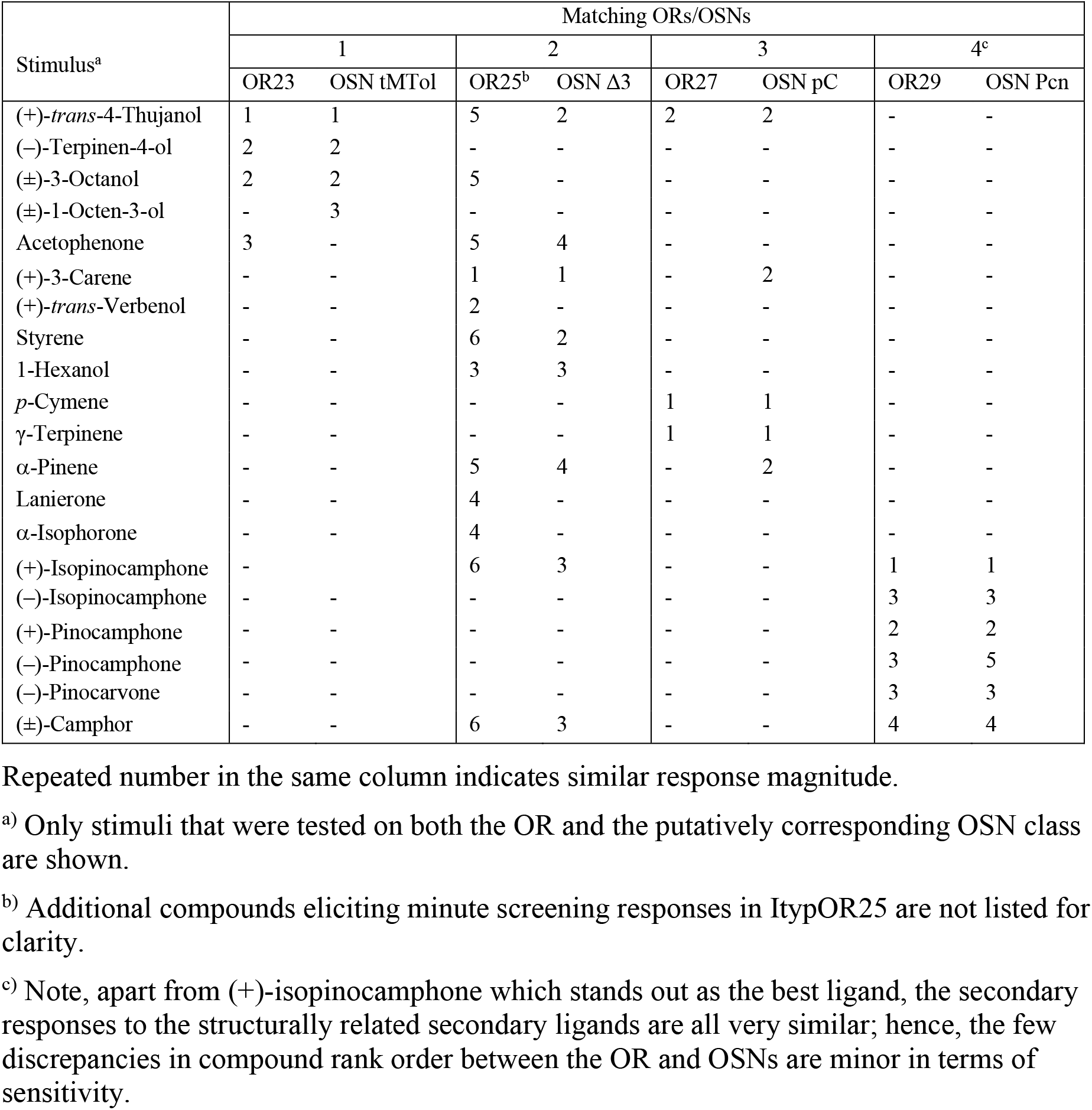
Comparing *in vitro* responses of odorant receptors (ORs) with OSN responses. Rank order (1 = best compound, 2 = second best, etc.) based on response magnitude (SSR data from Tømmerås, 1985; Andersson et al., 2009; Schiebe et al., 2019; Kandasamy, 2019). Normalized responses for ItypORs in oocytes.

**Fig. 2.**
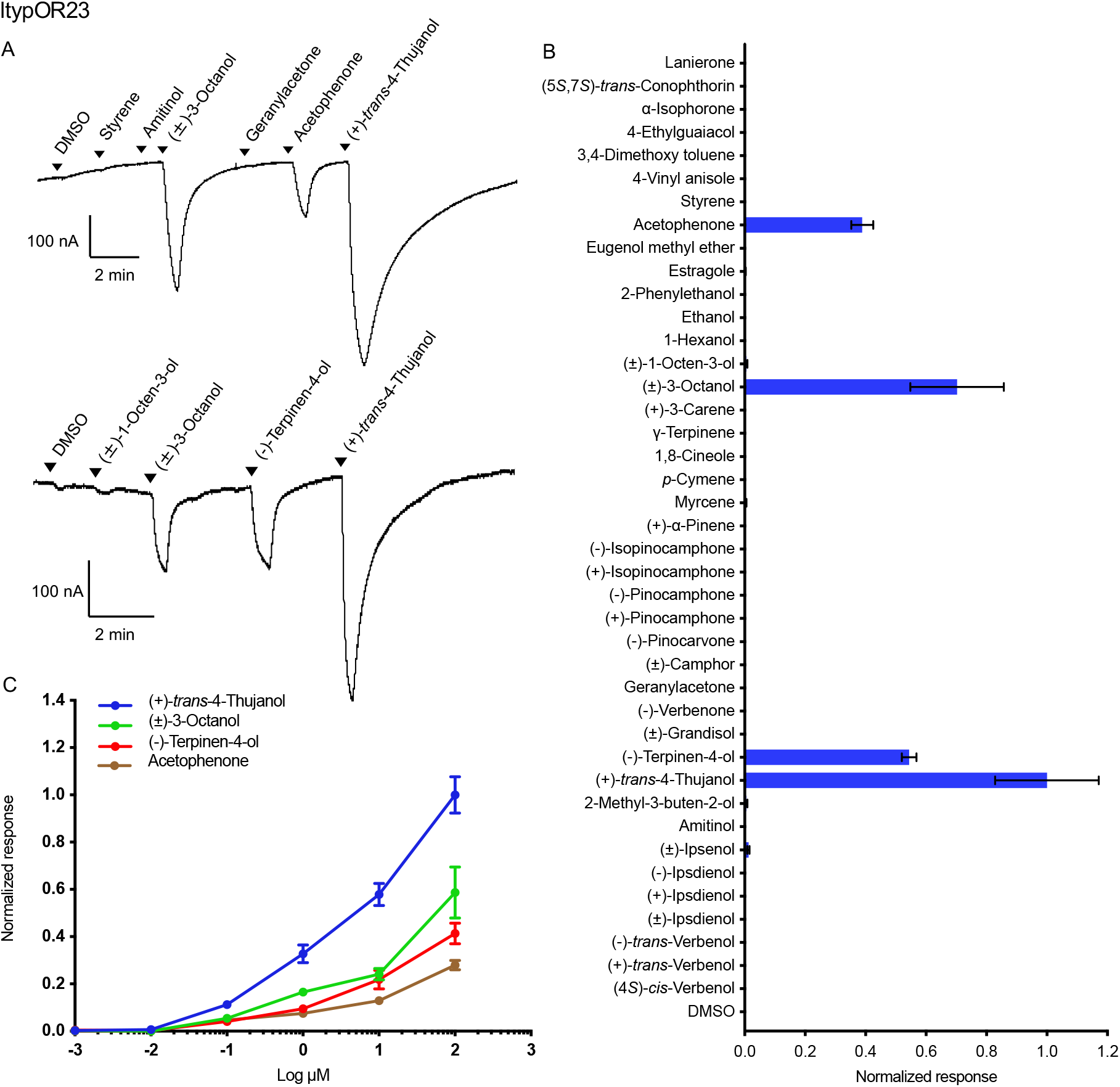
Responses of ItypOR23 in oocytes. (A) Representative current traces of oocytes upon successive exposures to 100 μM stimuli. Each compound was applied at the time indicated by the arrowheads for 20 s. Upper and lower traces include different sets of test stimuli. (B) Response profile of ItypOR23 to the full odor panel. Values were normalized based on the average response to the primary ligand of ItypOR23 (n ≥ 4). (C) Dose-dependent responses of oocytes expressing ItypOR23/Orco to active ligands. Values were normalized based on the average response of ItypOR23 to the most active compound at 100 μM (n ≥ 4 for each ligand). Error bars indicate the SE.

In the screening experiment, oocytes co-expressing ItypOR29 and ItypOrco responded most strongly to (+)-isopinocamphone, followed by rather strong secondary responses to the structurally similar ketones (+)-pinocamphone, (–)-pinocarvone, (–)-pinocamphone, (–)-isopinocamphone, and (±)-camphor (Fig. 3A and B). As a result of the thermodynamic equilibrium, both the synthesized pinocamphone enantiomers contained 16-19% impurities of the corresponding isopinocamphone enantiomers (Supplementary Table S1), and these may have influenced the responses the pinocamphones. Several additional compounds elicited weaker responses in the screening experiment (Fig. 3B). The dose-response assays confirmed that (+)-isopinocamphone is the primary ligand for this OR, although the responses elicited by the secondary ligands were also comparatively high and similar among the tested compounds (Fig. 3C). The responses of ItypOR29 are highly similar to the responses of the putatively corresponding OSN class (named OSN class Pcn; Schiebe et al., 2019; Kandasamy, 2019), with only minor discrepancies in the rank order among the secondary ligands, which are all similarly active in both systems (Table 1).

**Fig. 3.**
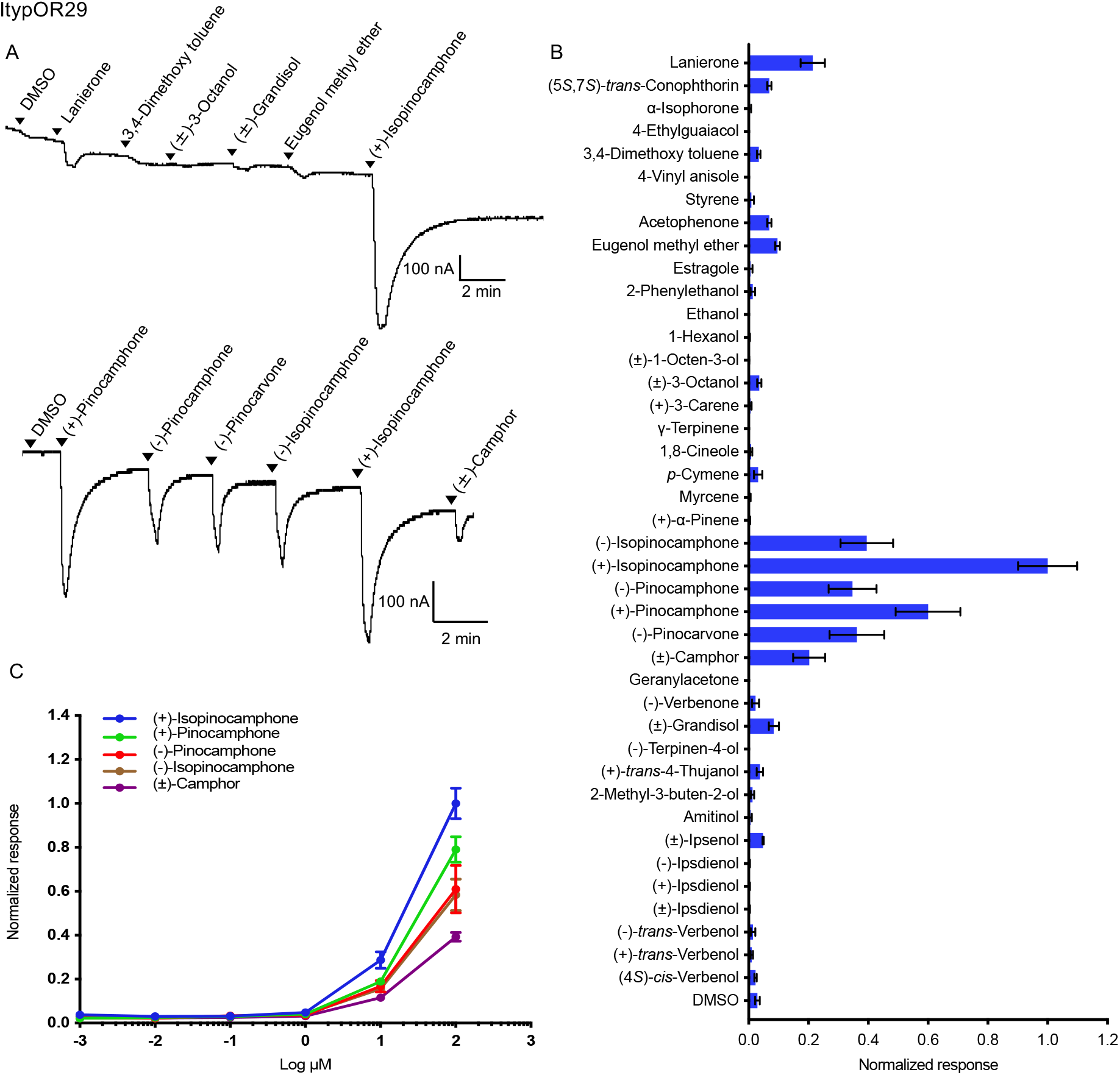
Responses of ItypOR29 in oocytes. (A) Representative current traces of oocytes upon successive exposures to 100 μM stimuli. Each compound was applied at the time indicated by the arrowheads for 20 s. Upper and lower traces include different sets of test stimuli. (B) Response profile of ItypOR29 to the full odor panel. Values were normalized based on the average response to the primary ligand of ItypOR29 (n ≥ 4). (C) Dose-dependent responses of oocytes expressing ItypOR29/Orco to active ligands. Values were normalized based on the average response of ItypOR29 to the most active compound at 100 μM (n ≥ 4 for each ligand). Error bars indicate the SE.

### ItypOR25 and ItypOR27 respond to host plant volatiles

*Xenopus* oocytes co-expressing ItypOR25 and ItypOrco responded to several compounds in the screening experiment. The strongest response was elicited by the conifer volatile (+)-3-carene followed by the beetle-produced compound (+)-*trans*-verbenol and the non-host plant volatile 1-hexanol. Weaker, yet clear, responses were elicited primarily by lanierone and α-isophorone, whereas responses to additional compounds, such as acetophenone, (±)-3-octanol and (+)-*trans*-4-thujanol, were minor (Fig. 4A and B). The dose-response experiments for ItypOR25 indicated a higher sensitivity to (+)-3-carene as compared to the secondary ligands (Fig. 4C). Previous SSR experiments characterized an OSN class primarily excited by 3-carene (Andersson et al., 2009; Kandasamy, 2019). In addition, several secondary compounds are also shared between this OSN class (named OSN class Δ3; Andersson et al., 2009) and ItypOR25, although the rank orders between them are partially different (Table 1).

**Fig. 4.**
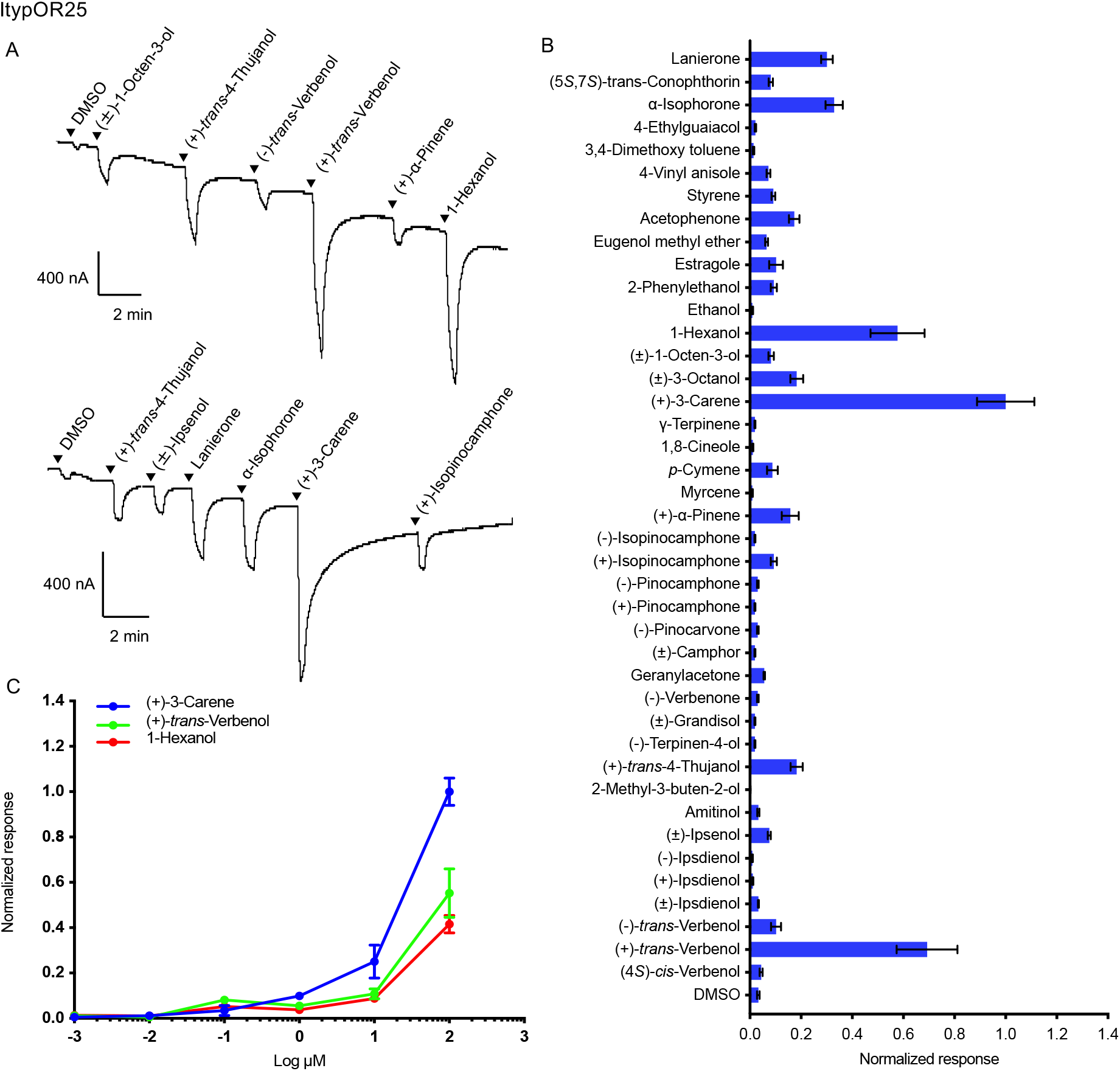
Responses of ItypOR25 in oocytes. (A) Representative current traces of oocytes upon successive exposures to 100 μM stimuli. Each compound was applied at the time indicated by the arrowheads for 20 s. Upper and lower traces include different sets of test stimuli. (B) Response profile of ItypOR25 to the full odor panel. Values were normalized based on the average response to the primary ligand of ItypOR25 (n ≥ 4). (C) Dose-dependent responses of oocytes expressing ItypOR25/Orco to active ligands. Values were normalized based on the average response of ItypOR25 to the most active compound at 100 μM (n ≥4 for each ligand). Error bars indicate the SE.

Oocytes co-expressing ItypOR27 and ItypOrco responded strongly and similarly to the two structurally similar host plant volatiles *p*-cymene and γ-terpinene in the screening assays (Fig. 5A and B). A weaker response was elicited by (+)-*trans*-4-thujanol. Dose-response experiments showed that ItypOR27 displayed a slightly higher sensitivity to *p*-cymene than to γ-terpinene, and a lower sensitivity to (+)-*trans*-4-thujanol (Fig. 5C). Similar to this OR, the OSN class originally reported to be highly specific for *p-*cymene (named OSN class pC; Andersson et al., 2009), was recently shown to respond just slightly less to γ-terpinene (Schiebe et al., 2019), and weaker responses to (+)-*trans*-4-thujanol and additional compounds have also been reported (Kandasamy, 2019). Hence, our data for ItypOR27 correspond very well with OSN data reported by Schiebe et al. (2019) (Table 1), whereas several of the weakly activating compounds reported by Kandasamy (2019), were inactive in our oocyte experiments.

**Fig. 5.**
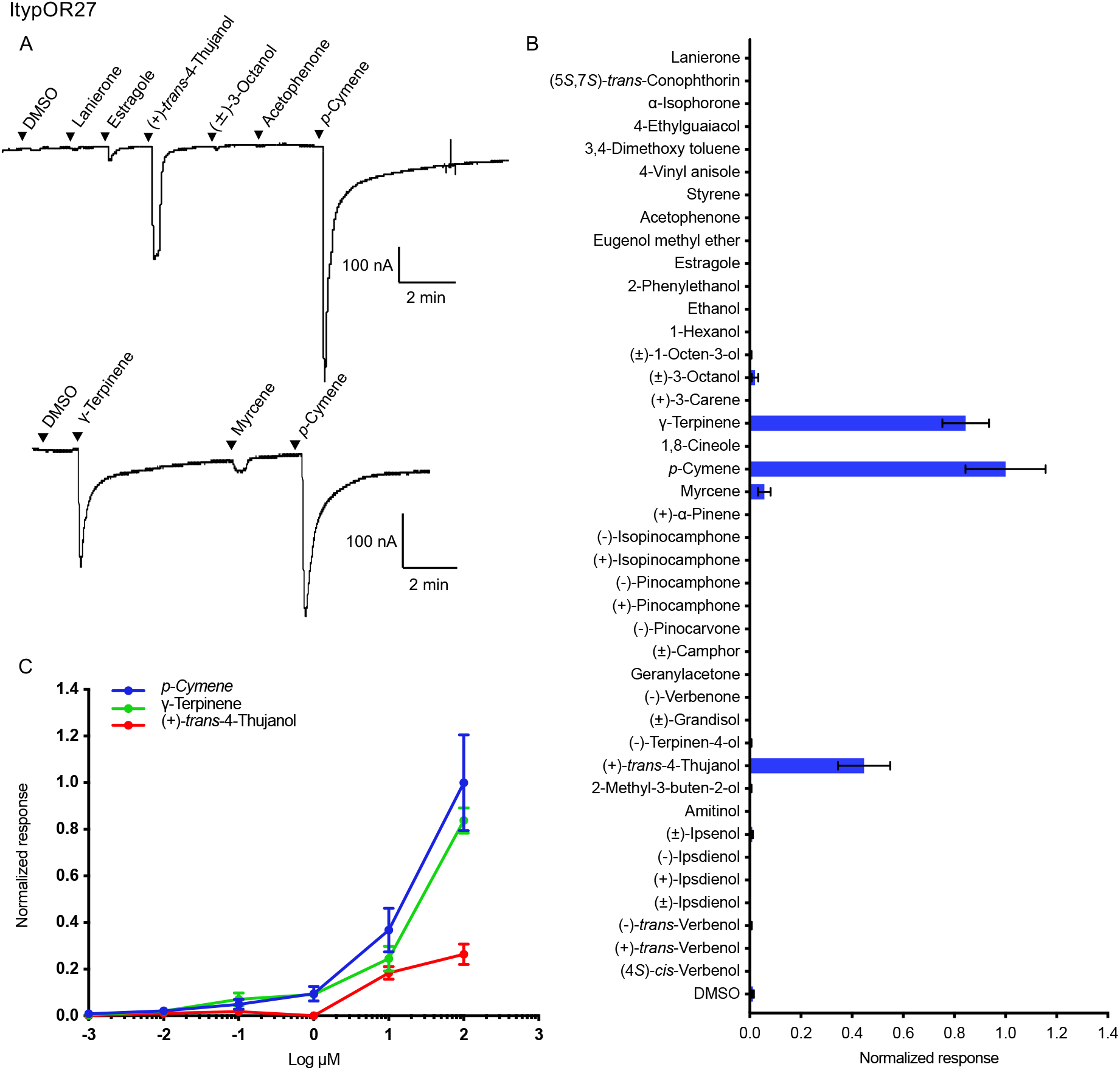
Responses of ItypOR27 in oocytes. (A) Representative current traces of oocytes upon successive exposures to 100 μM stimuli. Each compound was applied at the time indicated by the arrowheads for 20 s. Upper and lower traces include different sets of test stimuli. (B) Response profile of ItypOR27 to the full odor panel. Values were normalized based on the average response to the primary ligand of ItypOR27 (n ≥ 4). (C) Dose-dependent responses of oocytes expressing ItypOR27/Orco to active ligands. Values were normalized based on the average response of ItypOR27 to the most active compound at 100 μM (n ≥ 3 for each ligand). Error bars indicate the SE.

### *ItypOR28 responds to a pheromone component of the sympatric* I. duplicatus

Oocytes co-expressing ItypOR28 and ItypOrco were activated by bark beetle pheromones, showing the strongest response to *E*-myrcenol in the screening experiment. *E*-myrcenol is used as an aggregation pheromone component by the competitor *I. duplicatus* (Byers et al., 1990). Secondary responses were elicited by the structurally related compounds ipsdienol and ipsenol (Fig. 6A and B). Dose-response experiments that included *E*-myrcenol, racemic ipsenol, racemic ipsdienol and its two pure enantiomers showed that the response threshold of ItypOR28 is the lowest for *E*-myrcenol, compared with that for both ipsdienol and ipsenol (Fig. 6C). An OSN responding to *E*-myrcenol in *I. typographus* has to our knowledge not been reported.

**Fig. 6.**
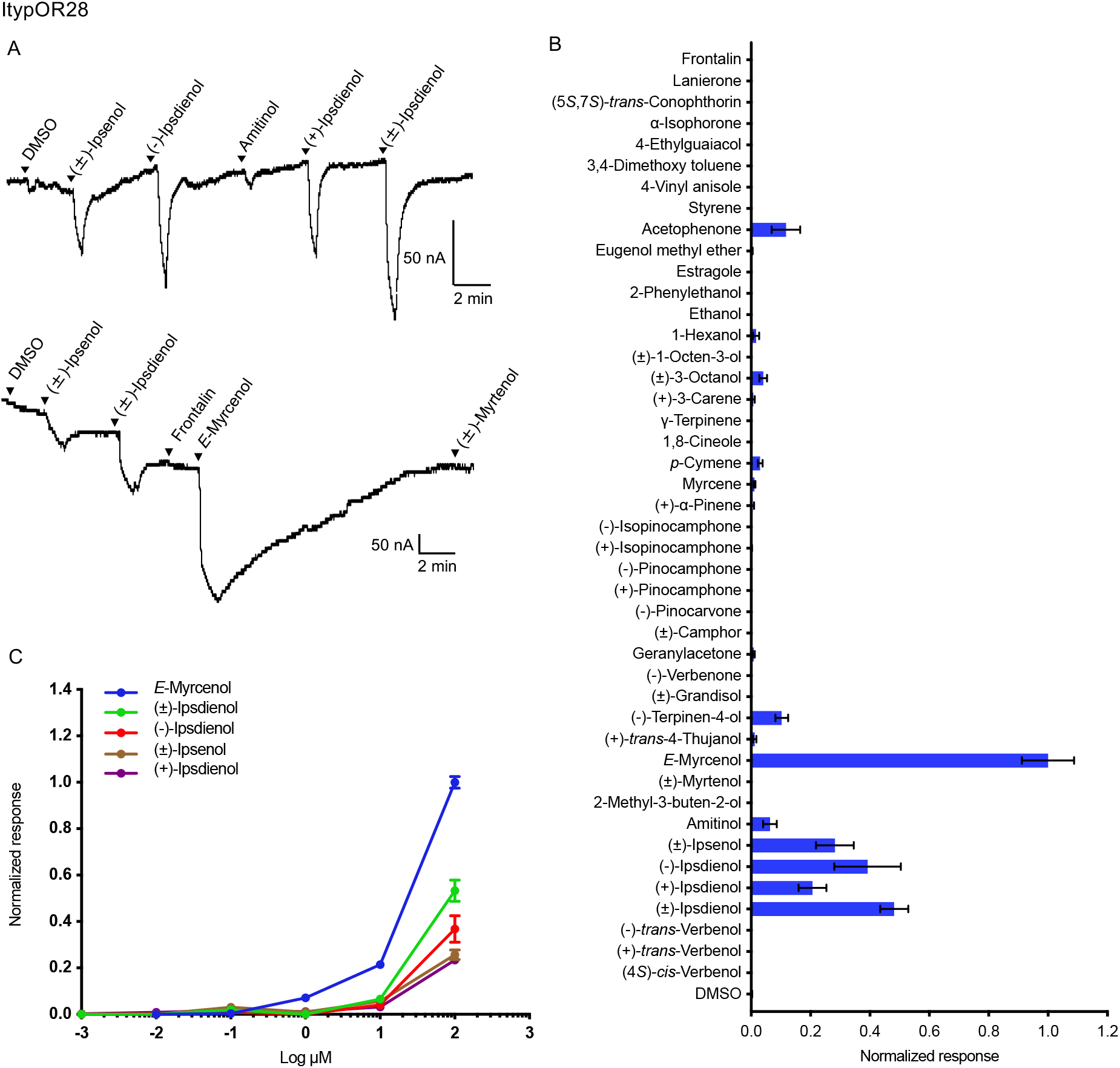
Responses of ItypOR28 in oocytes. (A) Representative current traces of oocytes upon successive exposures to 100 μM stimuli. Each compound was applied at the time indicated by the arrowheads for 20 s. Upper and lower traces include different sets of test stimuli. (B) Response profile of ItypOR28 to the full odor panel. Values were normalized based on the average response to the primary ligand of ItypOR28 (n ≥ 4). (C) Dose-dependent responses of oocytes expressing ItypOR28/Orco to active ligands. Values were normalized based on the average response of ItypOR28 to the most active compound at 100 μM (n ≥ 4 for each ligand). Error bars indicate the SE.

## Discussion

### *Functional evolution of the* I. typographus *OR-radiation*

A recent phylogenetic analysis (Yuvaraj et al., 2021) that included the 73 antennally expressed ORs of *I. typographus* along with the ORs from the mountain pine beetle *D. ponderosae* (Andersson et al., 2019) and the Asian longhorn beetle *Anoplophora glabripennis* (Cerambycidae) (McKenna et al., 2016) revealed a highly supported clade formed by seven ItypORs. Two of these receptors (ItypOR46 and ItypOR49) were shown to selectively respond to the pheromone compounds (*S*)-(–)-ipsenol (ItypOR46) and (*R*)-(–)-ipsdienol (ItypOR49) (Yuvaraj et al., 2021). Here, we obtained functional data for the five remaining ItypORs from this clade. Collectively, our results show that this clade contains two ORs specifically tuned to monoterpene hydrocarbons produced by the host tree (ItypOR25 and ItypOR27), three ORs tuned to oxygenated monoterpenoids produced by con-or heterospecific bark beetles (i.e., pheromone compounds; ItypOR28, ItypOR46, and ItypOR49), and two ORs responding to volatiles produced by the symbiotic fungi of *I. typographus* (ItypOR23 and ItypOR29). Similar to previous suggestions (Mitchell et al., 2012; Mitchell and Andersson 2020; Yuvaraj et al., 2021), these findings strengthen the idea that pheromone receptors from different beetle species do not cluster in specific clades like they tend to do in Lepidoptera where the majority of the characterized pheromone receptors (PRs) are found in the so called classical PR clade (Zhang and Löfstedt, 2015); instead beetle pheromone receptors are scattered in the OR phylogeny, and OR clades include receptors detecting compounds from various ecological sources. This may not be surprising because most bark beetle pheromones are oxygenated monoterpenoids (sometimes produced through metabolism of host monoterpenes) and hence structurally similar to common conifer host tree compounds (and to several fungal-derived compounds), contrasting the situation in moths where most pheromone compounds are structurally different from plant volatiles. In addition, host-and mate localization in bark beetles is an integrated process involving pheromones, host and non-host odors, with aggregation pheromones mediating sex-attraction, aggregation behavior, and successful localization and infestation of host trees providing food.

While our functional data suggest a clade in which all ItypORs detect monoterpenes or monoterpenoids and are under strong purifying selection, our findings also bring up questions of how odor specificity may evolve in OR-lineage radiations. Whereas the pheromone compounds ipsenol, ipsdienol, and *E*-myrcenol are similar acyclic monoterpenols with a myrcene backbone, the other ligands for the ORs in this clade include monoterpene hydrocarbons and ketones as well as cyclic structures (Fig. 1D). The splits between the seven ORs appear to be rather old, as indicated by the branch lengths (Fig. 1A) and low amino acid identities (Fig. 1C), and this appears to have provided sufficient time for significant specificity shifts (neofunctionalization) to evolve. Purifying selection against deleterious mutations then appears to have dominated to retain the OR responses to these ecologically important compounds.

Previous work (Tømmerås et al., 1984; Andersson et al., 2009) identified OSNs primarily responding to e.g. (*S*)-(+)-ipsdienol and amitinol, which are both structurally related to the primary ligands (*S*)-(–)-ipsenol, (*R*)-(–)-ipsdienol, and *E*-myrcenol of ORs characterized here. Unexpectedly, the ORs detecting these two similar compounds are not found within this receptor lineage, and hence must have evolved their specificities independently. The same holds true for the several OSN classes specifically tuned to monoterpene hydrocarbons or oxygenated monoterpenoids (Andersson et al., 2009; Schiebe et al., 2019; Kandasamy, 2019), of which only four putatively corresponding ORs are localized in this clade.

Across several species of Lepidoptera, the structures of key ligands for characterized ORs follow the OR phylogeny at a broader scale, with ligands detected by relatively closely related ORs sharing similar molecular features (de Fouchier et al., 2017; Guo et al., 2021). But similar to our findings, there is also some variation within OR clades with respect to e.g., functional groups and unsaturation among ligands detected by related ORs. For example, the recently identified “novel” PR clade in Lepidoptera harbors receptors responsive to structural isomers of the ligands for the receptors in classical PR clade (Zhang and Löfstedt, 2015), although the two clades are distant (Bastin-Héline et al., 2019). It was suggested that Lepidoptera PRs have evolved independently multiple times, although most of them reside in the classical PR clade.

To our knowledge, our study represents a rare example where all ORs in a species-specific OR-radiation comprising as many as seven receptors have been functionally characterized. Hence, large-scale functional characterization of the OR repertoires of *I. typographus* and additional species is required to further our understanding of the functional evolution of the insect OR gene family.

### ItypOR23 and ItypOR29 detect volatiles mainly produced by fungal symbionts

Like other bark beetles, *I. typographus* is associated with a suite of symbiotic ophiostomatoid fungi, which beetles inoculate into the inner bark of attacked trees. These fungi may serve as food for maturing beetles (Kandasamy et al., 2019) and they also metabolize spruce defense compounds (Kandasamy, 2019). In laboratory bioassays, beetles actively seek out fungal-colonized media using olfactory cues and are able to distinguish different fungal species by their different odor profiles using OSNs dedicated for fungal volatiles (Kandasamy et al., 2019). Here, we characterized two receptors, ItypOR23 and ItypOR29, responding to the fungal-derived volatiles (+)-*trans-*4-thujanol and (+)-isopinocamphone, respectively. Both of these ORs showed high response specificities to only a few structurally related compounds. The OR response profiles matched very well with those from the characterized OSNs, where the tMTol class is highly specific for (+)-*trans-*4-thujanol and the Pcn class selective for the highly similar monoterpene ketones that activate ItypOR29 (Table 1; Schiebe et al., 2019; Kandasamy, 2019).

It was recently shown that the fungi of *I. typographus* transform the host compounds (–)-β-pinene, (–)-α-pinene and (–)-bornyl acetate to oxygenated monoterpenoids, including the primary and secondary ligands of both ItypOR23 and ItypOR29 (Kandasamy, 2019). In lab bioassays, (+)-*trans*-4-thujanol and camphor were attractive to adult beetles at a relatively low dose (100 µg), however, (+)-*trans*-4-thujanol was repellent at higher doses (200 µg to 1 mg) (Blažytė-Ĉereškienė et al., 2016; D. Kandasamy, M. N. Andersson, A. Hammerbacher, J. Gershenzon et al., unpublished data). This suggests concentration-dependent effects of these compounds and that changes in their emission over time may be used by new incoming bark beetles to evaluate the colonization status of the host tree. In addition to production by fungi, these oxygenated monoterpenoids are also released from spruce trees, with amounts decreasing with tree age and increasing with tree stress (Blažytė-Čereškienė et al., 2016; Schiebe et al., 2019). Hence, these compounds of multiple ecological origins may be also used by *I. typographus* to assess the physiological status of host trees.

### ItypOR25 and ItypOR27 are tuned to host plant volatiles

ItypOR25 and OR27 were most sensitive to two host monoterpenes, (+)-3-carene and *p*-cymene, respectively, and especially ItypOR27 demonstrated a high specificity. These two ORs group together within the analyzed OR clade and present the most recent split among the seven ORs, which is in line with a tuning shift from detecting oxygenated monoterpenoids to monoterpene hydrocarbons. Despite this shared characteristic, the response profiles of the two ORs are widely different suggesting that neofunctionalization has been favored also after their presumed duplication (Rastogi and Liberles, 2005; Andersson et al., 2015). Whereas the response profile of ItypOR27 matches very well with that of the pC OSN class, ItypOR25 presented a worse match with a broader response profile and partly different rank orders among the secondary compounds compared to the putatively corresponding Δ3 OSN class (Table 1; Andersson et al., 2009; Schiebe et al., 2019). It remains unknown whether this discrepancy is due to the artificial oocyte environment or whether ItypOR25 is not the receptor of the Δ3 OSN class. However, differences in OR responses between different *in vitro* systems and between these systems and the OSNs have repeatedly been found in previous studies, including for ItypOR46 and ItypOR49 and corresponding OSN classes (Yuvaraj et al., 2017; Yuvaraj et al., 2021; Hou et al., 2020). Because of this system-dependency and since no other OSN in *I. typographus* has shown a primary response to 3-carene (Andersson et al., 2009; Schiebe et al., 2019; Kandasamy, 2019; Kandasamy et al., 2019), it appears likely that ItypOR25 is indeed the receptor that corresponds to the Δ3 OSN class.

The roles of host monoterpenes in host selection by *I. typographus* is poorly understood and likely context-dependent. While attraction to host monoterpenes alone has not been demonstrated, the major host compounds α-pinene and β-pinene can enhance the response of the bark beetle to the aggregation pheromone (Rudinsky et al., 1972; Hulcr et al., 2006; Erbilgin et al., 2007). In contrast, less abundant host compounds may reduce the pheromone attraction. For example, 1,8-cineole is more abundant in resistant trees and upregulated during an attack, and this compound has a strong antagonistic effect on pheromone attraction (Andersson et al., 2010; Schiebe et al., 2012; Binyameen et al., 2014). Similarly, *p*-cymene is increased in heavily attacked trees, and it reduced pheromone trap catch by 50% (Andersson et al., 2010). Hence, increased amounts of inducible toxic host compounds may be used by bark beetles as cues to evaluate host defense potential and/or breeding density, and may therefore represent general ‘host unsuitability’ signals (Erbilgin and Raffa, 2000; Seybold et al., 2006; Andersson et al., 2010) with context-and species-dependent effects (Saint-Germain et al., 2007; Raffa et al., 2016). To our knowledge, behavioral effects of 3-carene have not been studied in *I. typographus*. However, the amount of 3-carene in Norway spruce increases after inoculation with the *I. typographus*-associated fungus *Endoconidiophora polonica*, and this compound has been linked to conifer resistance and susceptibility to insects and fungi (Zhao et al., 2010, and references therein). In addition, 3-carene is also abundant in pine trees, including sympatric Scots pine (*Pinus sylvestris*) and it may hence be used as a host suitability cue, similar to *p*-cymene and 1,8-cineole.

### ItypOR28, ItypOR46, and ItypOR49 detect bark beetle pheromones

We characterized one bark beetle pheromone receptor (ItypOR28) primarily responding to *E*-myrcenol. Hence, together with the previously characterized ItypOR46 ((*S*)-(–)*-*ipsenol) and ItypOR49 ((*R*)-(–)-ipsdienol) (Yuvaraj et al., 2021), the targeted clade houses three pheromone receptors. The ipsenol and ipsdienol enantiomers are produced by male *I. typographus* during the later attack phases (2-6 days after boring has been initiated), and at least ipsenol inhibits the attraction to the aggregation pheromone, possibly to avoid intraspecific competition (Francke et al., 1980; Birgersson et al., 1984; Schlyter et al., 1989). *E-*myrcenol together with both enantiomers of ipsdienol comprise the aggregation pheromone of *I. duplicatus*, the largest competitor of *I. typographus* (Byers et al., 1990), and *E-*myrcenol inhibits the attraction of *I. typographus* to its aggregation pheromone (Schlyter et al., 1992). Consistent with a general high specificity of insect pheromone receptors (Andersson et al., 2015), ItypOR28 only responded secondarily to the structurally related ipsdienol and ipsenol, and narrow tunings were also reported for ItypOR46 and ItypOR49 (Yuvaraj et al., 2021). In contrast to the four other ItypORs characterized here, no OSN class has been reported to be specific for *E-*myrcenol despite large screening efforts that included this compound (Andersson et al., 2009). Our results from ItypOR28 suggest the existence of such an OSN class, and it is possible that these OSNs have been missed because some OSN classes were recently shown to have a highly restricted spatial distribution on the *I. typographus* antenna (Kandasamy et al., 2019).

### Concluding remarks

We determined the functions of five ORs from a species-specific clade formed by seven receptors in *I. typographus* using the *Xenopus* oocyte system. Together with the two pheromone receptors previously characterized in HEK cells and oocytes (Yuvaraj et al., 2021), this clade contains seven ORs which respond to monoterpenes or monoterpenoids, indicating the biological importance of such compounds to *I. typographus*. Three ORs respond to bark beetle pheromones, two ORs to host plant volatiles, and two ORs to (mainly) fungal-derived compounds. The structural similarities especially among the three pheromonal ligands detected by ItypOR28, ItypOR46, and ItypOR49 implies that the ORs in this species-specific radiation have evolved specificities for chemically similar compounds. In addition, the variation in the structures of the key ligands for the seven receptors and their various ecological origins suggest that neofunctionalization has been promoted in the evolution of this OR clade after which purifying selection has dominated to retain the new OR responses. Finally, our results indicate that the pheromone receptors of beetles do not form specific pheromone receptor clades as seen in Lepidoptera (Yuvaraj et al., 2018). Instead, beetle pheromone receptors are dispersed across the OR phylogeny, and OR-lineage radiations within species contain receptors that detect compounds of different ecological origins.

## Materials and methods

### Chemicals

Compounds tested in this study were from commercial sources, synthesized by the Unelius laboratory or obtained from colleagues as gifts (Table S1). The test compounds included beetle-produced pheromone compounds, and volatiles produced by associated fungi, conifer host and angiosperm non-host trees, including the key ligands for all known OSN classes of *I. typographus* as well as several secondary ligands for these neurons (Tömmerås, 1985; Andersson et al., 2009; Schiebe et al., 2019; Kandasamy, 2019; Kandasamy et al., 2019). Stock solutions for *Xenopus* oocyte recordings were prepared by diluting each compound to 100 mM in dimethyl sulfoxide (DMSO), which were stored at -20°C. Before each experiment, the stock solutions were diluted to desired concentration in Ringer’s buffer (96 mM NaCl, 2 mM KCl, 5mM MgCl2, 0.8 mM CaCl2, 5 mM HEPES, pH 7.6) with the final stimuli containing 0.1% DMSO. Ringer’s buffer containing 0.1% DMSO was used as negative control. Compounds were initially screened for receptor activity at a concentration of 100 μM, and the active compounds were subsequently tested in dose-response experiments using six concentrations at 10-fold increments ranging from 1 nM to 100 µM. An odor panel of 32 compounds was initially tested on all five ORs (ItypOR23, 25, 27, 28, and 29) (Table S1). After obtaining the first screening responses to this odor panel, additional OR-specific compounds (putative secondary ligands) were selected for testing based on previous data from putatively matching OSN types, which facilitated our comparisons of OR responses with those of previously characterized OSN classes (Table S2). For consistency, these nine additional compounds were also tested on the other non-target ORs over two replications (showing insignificant activity). For ItypOR28, the initial screening experiment indicated only minor and rather unspecific responses to some test compounds. Hence, we hypothesized that this OR may be tuned to a structurally related compound, and therefore also tested frontalin, *E*-myrcenol and myrtenol specifically on this OR (Table S3). Chemical structures presented in Fig. 1D were drawn using ChemDraw Professional 17.0.

### Vector construction and cRNA synthesis

Full length sequences of ItypORs and ItypOrco were amplified by PCR using gene specific primers containing flanking 5’ Kozak sequence (‘GCCACC’) and 5’ and 3’ recognition sites for restriction enzymes (Table S4). The expression vectors pcDNA5/TO containing the five ItypORs (ItypOR23, 25, 27, 28, and 29; amino acid sequences are available in the Supplementary Material) and pcDNA4/TO containing ItypOrco that were used in previous HEK293 cell assays (Yuvaraj et al., 2021) were used as templates for sub-cloning. Sequences of the five ItypORs have been deposited in GenBank under the accession numbers MW556722-MW556726. The PCR products were analyzed on 1% TAE agarose gels, and then purified using the GeneJET Gel Extraction Kit (Thermo Fisher Scientific™). The purified fragments were ligated into the oocyte expression vector pCS2+ using T4 ligase (Thermo Fisher Scientific) overnight at 4 °C and then transformed into TOP10 competent cells (Thermo Fisher Scientific). Positive colonies that were confirmed by colony PCR were grown in LB broth overnight with ampicillin, and the plasmids were then extracted using the GeneJET plasmid miniprep kit (Thermo Fisher Scientific). Plasmids were Sanger sequenced on a capillary 3130xL Genetic Analyzer (Thermo Fisher Scientific, Waltham, MA, USA) using Sp6 and EBV-rev primers at the sequencing facility at the Department of Biology, Lund University. Large quantities of purified plasmids containing ItypORs and ItypOrco with correct sequences were obtained using the PureLinkTM HiPure Plasmid Filter Midiprep Kit (Thermo Fisher Scientific). cRNAs of ItypORs and ItypOrco were synthesized from NotI (Promega) linearized recombinant pCS2+ plasmids using the mMESSAGE mMACHINE SP6 transcription kit (Thermo Fisher Scientific).

### Microinjection and two-electrode voltage clamp recordings

Each of the five ItypORs were co-expressed with ItypOrco in *Xenopus laevis* oocytes, and two-electrode voltage clamp recordings were performed following previously described protocols (Zhang et al., 2010; Zhang and Löfstedt, 2013). In brief, oocytes were surgically collected from *X. laevis* females (frogs were purchased from University of Portsmouth, UK) and treated with 1.5 mg/mL collagenase (Sigma-Aldrich Co., St. Louis, MO, USA) in oocyte Ringer 2 buffer (82.5 mM NaCl, 2 mM KCl, 1mM MgCl2, 5mM HEPES, pH 7.5) at room temperature for 15-18 min. Mature healthy oocytes (stage V-VII) were co-injected with 40 ng each of candidate OR and Orco cRNAs, then incubated for 3-7 days at 18 °C in Ringer’s buffer containing sodium pyruvate (550 mg/L) and gentamicin (100 mg/L). Odorant-induced whole-cell inward currents were recorded from injected oocytes in good condition using a two-electrode voltage clamp with a TEC-03BF amplifier at the holding potential of -80 mV. A computer-controlled perfusion system was employed to apply the tested compounds and Ringer’s buffer to the oocyte chamber. Oocytes were exposed to compounds at a flow rate of 2 mL/min for 20 s with extensive washing with Ringer’s buffer at a flow rate of 4 mL/min between stimulations to recover the current to baseline. Data were collected and analyzed using the Cellworks software (NPI Electronic GmbH, Tamm, Germany), and the responses were normalized by calculating the relative response to each active compound in relation to the average response to the most active compound.

### Evolutionary analysis of ORs

Amino acid sequences of select Group 7 ORs (Mitchell et al., 2020) from *I. typographus* and *D. ponderosae* were aligned using MAFFT (v7.450; Katoh et al., 2002), and an unrooted maximum likelihood tree was constructed using FastTree 2.1.11 (Price et al., 2010) implemented in Geneious Prime (2020.0.5) software (Biomatters Ltd., Auckland, New Zealand). Local node support values were calculated using the Shimodaira-Hasegawa (SH) test implemented in FastTree. The nonsynonymous (dN) to synonymous (dS) substitution rate (ω) was estimated by the maximum likelihood method (Anisimova et al., 2001) using the Codeml program in the PAML 4.6 package (Yang, 1997).

## Acknowledgements

We are grateful to Dineshkumar Kandasamy, Suresh Ganji, Erika Wallin, Blanka Kalinová, Fredrik Schlyter, Anna Jirošová, and Aleš Machara for providing test compounds. We thank the Chinese Scholarship Council (CSC) for finical support of Xiaoqing Hou’s PhD study. This work was also supported by grants from the Swedish Research Councils FORMAS (grants #217-2014-689 and #2018-01444 to M.N.A.) and VR (#2017-03804 to C.L.), the Crafoord foundation (M.N.A.), the Carl Trygger foundation (M.N.A.), and the Royal Physiographic Society in Lund (R.E.R. and M.N.A.). The study was carried out as part of the Max Planck Centre for Next Generation Chemical Ecology (nGICE).

## Data availability

All oocyte data are presented in the main figures of the paper. Sequences of the characterized ItypORs have been deposited in Genbank (accession numbers MW556722-MW556726).

