## Supplementary Material for "Functional evolution of a bark beetle odorant receptor clade detecting monoterpenoids of different ecological origins"

**Table S1.** List of compounds initially tested on all five ORs, with purity, source and examples of biological origin.

| Class | Compound | Purity (%) | Source* | Biological origin |
|---|---|---|---|---|
| Oxygenated monoterpenoids homoterpenoids, and hemiterpenes | (1*S*,4*S*)-*cis*-Verbenol | 95 | Borregaard | Beetle |
| | (1*R*,4*R*)-(+)-*trans*-Verbenol | 92 | SCM | Beetle |
| | (1*S*,4*S*)-(–)-*trans*-Verbenol | 97 | SciTech Ltd., Prague | Beetle |
| | (1*S*)-(–)-*cis*-Verbenone | >99 | Fluka | Beetle, fungi |
| | (±)-Ipsdienol | 94 | Bedoukian | Beetle |
| | (±)-Ipsenol | 95 | Synergy Semiochemicals | Beetle |
| | Amitinol | 91 | R. U. | Beetle |
| | (+)-*trans*-4-Thujanol | 97 | Sigma-Aldrich | Host, fungi |
| | (±)-Grandisol (grandlure I) | 95 | Bedoukian (E. W.) | Beetle |
| | Geranylacetone | >99 | Fluka | Non-host, fungi |
| | (+)-Isopinocamphone | >99 | R. U. | Host, fungi |
| | 1,8-Cineole | 99 | Aldrich | Host |
| | α-Isophorone | >99 | Acros | ** |
| | Lanierone | >99 | Synergy Semiochemicals | Beetle |
| | 2-Methyl-3-buten-2-ol | >99 | Acros | Beetle, fungi |
| Monoterpenes | (+)-α-Pinene | 98 | Janssen Chimica | Host |
| | Myrcene | 95 | Sigma-Aldrich | Host |
| | *p*-Cymene | >99 | Acros | Host |
| | (+)-3-Carene | 99 | Aldrich | Host |
| Aliphatic alcohols and spiroacetals | (±)-3-Octanol | 97 | Sigma-Aldrich | Non-host, fungi |
| | (±)-1-Octen-3-ol | 98 | Janssen Chimica | Non-host, fungi |
| | 1-Hexanol | >99 | Fluka | Non-host, fungi |
| | Ethanol | 99.5 | Solveco | Host |
| | (5*S*,7*S*)-*trans*-Conophthorin | 94 | W. F. | Non-host, fungi |
| Aromatic compounds | 2-Phenylethanol | >99 | Sigma | Beetle, fungi |
| | Acetophenone | 99 | Acros | Beetle, fungi |
| | Styrene | >99 | Fluka | Fungi |
| | 4-Vinylanisole | 97 | Aldrich | Fungi |
| | Estragole (4-allylanisole) | >99 | Aldrich | Host, fungi |
| | 3,4-Dimethoxytoluene | 98 | Givaudan-Roure | Host, fungi |
| | Eugenol methyl ether | >99 | Fluka | Host, fungi |
| | 4-Ethylguaiacol | 98 | Sigma-Aldrich | Fungi |

*R. U. = synthesized by Rikard Unelius (Linnaeus University, Kalmar, Sweden); W. F. = gift from Wittko Francke (University of Hamburg, Germany). **: The biological source of α-isophorone is unknown. Female *Ips typographus* produce tiny amounts of β-isophorone which is unstable, and we found that α-isophorone was more active than β-isophorone in SSR study (Kandasamy et al., in prep), so we used α-isophorone in this study.

**Table S2.** Additional OR-specific compounds, with purity, source, examples of biological origin and target ORs. For consistency, these compounds were also tested on non-target ORs showing no (or insignificant) activity.

| Class | | Purity (%)* | Source** | Biological origin | Target OR |
|---|---|---|---|---|---|
| Oxygenated monoterpenoids | (*S*)-(+)-Ipsdienol | 99 (98% ee) | A. M. | Beetle | ItypOR28 |
| | (*R*)-(–)-Ipsdienol | 99 (98% ee) | A. M. | Beetle | ItypOR28 |
| | (–)-Terpinene-4-ol | 99 | Acros | Host, fungi | ItypOR23 |
| | (±)-Camphor | 97 | Aldrich | Host, fungi | ItypOR29 |
| | (–)-Pinocarvone | 99 | D. K. | Beetle, fungi | ItypOR29 |
| | (+)-Pinocamphone | 84 (16% IPC) | R. U. | Host, fungi | ItypOR29 |
| | (–)-Pinocamphone | 81 (19% IPC) | R. U. | Host, fungi | ItypOR29 |
| | (–)-Isopinocamphone | 99 | R. U. | Host, fungi | ItypOR29 |
| Monoterpene | γ-Terpinene | 97 | Aldrich | Host | ItypOR27 |

Abbrevations: ee = Enantiomeric excess; IPC= isopinocamphone.

** R. U. = synthesized by Rikard Unelius (Linnaeus University, Kalmar, Sweden);  A. M. = synthesized by Aleš Machara (Academy of Sciences of the Czech Republic, Prague);

D. K. = gift from Dineshkumar Kandasamy (Max Planck Institute for Chemical Ecology, Jena, Germany).

**Table S3.** Three compounds only tested on ItypOR28, with purity, source, and examples of biological origin.

| Class | Compound | Purity (%) | Source* | Biological origin |
|---|---|---|---|---|
| Oxygenated monoterpenoids | *E*-Myrcenol | >99 | Fytofarm | Beetle |
| | (±)-Myrtenol | 96 | G. B. | Beetle, fungi |
| Bicyclic acetal | (±)-Frontalin | >99 | Synergy Semiochemicals | Beetle |

* G. B. = gift from Gunnar Bergström (University of Gothenburg, Sweden).

**Table S4.** Primers used in this study.

| Genes | Primer sequence (5’-3’) |
|---|---|
| ItypOR23_F | CGCGGATCCGCCACCATGGCCGTGTATCCAAAATCAG |
| ItypOR23_R | GCTCTAGATTAAGTTCTTTTGTATGCTAGAGTAATATAAGAAT |
| ItypOR25_F | CGCGGATCCGCCACC ATGAAGATTTACCCTGACACAAAGT |
| ItypOR25_R | GCTCTAGA CTATCGAAACATGATAGTAATATAGGTGTAGG |
| ItypOR27_F | CGCGGATCCGCCACC ATGAGAGTGTATCCGGACATAGAA |
| ItypOR27_R | GCTCTAGATTAGTTGTTTCGGACTACCACG |
| ItypOR28_F | CGCGGATCCGCCACC ATGGGATTGTATCCAGCAAGTAGA |
| ItypOR28_R | GCTCTAGATTAATTTCTAAGAATTACCGATATGTAGGTGT |
| ItypOR29_F | CGCGGATCCGCCACCATGGCTGCTTATCCACAATGC |
| ItypOR29_R | GCTCTAGACTATTGCCTAAAAATAATACTCACATAGGAATAA |
| Sp6 | ATTTAG GTGACACTATAG |
| EBV-rev | GTGGTTTGTCCAAACTCATC |

### Amino acid sequences of the five ItypORs.

>ItypOR23
MAVYPKSEHLKVPAIYCSTIGIFPWKFMFQDNKNLQTIYRCYSIVMLAWCIGFVVTDYIQLVILLTSKTLDMQEISFN
TCITLLFTCIGLRAVIVYFSPNSANLIQSIIDSEKVTYLDDAECMKLEKEHLRSVRLISHCYFIFIIFSTTSRCVYFFSKE
PDFIQNGNETEIVKEHMLSIWFPFNQEKYYLTVYNIELLDSFLGTFFVAYVDIYTFNMISYPKGQLKKLQHIMKHFH
NYKAKYSSETNEENDFIVFKDLVQRHKQIIQHINAFNELMEFVAIFEFVQSSAQIACGLTQTSLENLTIGSFLFVMSF
LISMLVRLFLYYYAANDVTVESTKLAQCIWESNWYEESQKIKLSMLMVIIRAQKPLIFKIGGFGTMSVQSIVTILKATY
SYITLAYKRT

>ItypOR25
MKIYPDTKFFDVTAKFGAIVGLYPWQFMFPDNNTCRQIYRWYSYIVLLSFIVLLLPMYVELIILLRNEETSKDELGSN
LSITIVFSSAGLRALFLRRGSNLINLIQNVMDEEKKQLFVDCKKVQLLEDKCLKVVRKLSYIYAVIVVVAASQKSVTA
LLQTPTSSTGTPSRDLIISAWFPFDKQEYYWQAYCIQIYHTIIGASYLSYMDIFMFNLLSYPIGQFKKLQFIIKNMEVQ
HYSYNDSENKSIDDGVRSIIERHQYIIQYVDFYNKSMGTFALFDFLQSSLQIATVLLQFSPTVGTIIFMLIFFALMLLR
LFLYYYTANEVSVQSEKVKMAVWESKWYEQPPKIKYALLRIMTRAAKPSKYIIGAFGGMSTYSIIQILKATYTYITIM
FR

>ItypOR27
MRVYPDIENFKITAIYSSTIGLFPWKFMFQDNQVLQQTYRYYSYFIYGSFVIFIITAYVELIIMLNGDVLKMDAICSNIC
LTLAFTCSALRATVMRVGPNLLKIIEQVMHAEKNPASIEDQTSFNLERKSIKTMRKLSHLYAVAITMIASSKCALAPF
EKGEIVHIGNTTIIDRPLIMSAWVPFNKNTHYWAAYIIQIYFAALGAWHVAYVDMFMFNMLGYPIGQLKKLHYYIKNI
TTLTRNDDSLEEFKNVIRQHQQIISYVKFYNDSMGTFAIFEFLQSSVQIASIFIQTSPSDMNLGQFGFIGGFFIGMLF
RLFLYYYTANEVMTESEKVGVSVWESDWYEQPTNLKMALLTVMMRGQRPLYYKIGGFGLMSVQSIVAILKATYT
YLTVVVRNN

>ItypOR28
MGLYPASRYFKNPIMWSSILGAFPWQMIFQENAKLQQVYRWYSNFMLTWYFGMVTTEYIQLYHILNANVIQMDE
VCENVCMSLVFTCTGLRVWVMRRTNGLSEIIQTVVDAEREADGLDDEKTRQYEDIHVKHMEKVSFIYAAFVFMSV
TNGCLATLYADTKSVIIGNSTIVEKPLIISTWFPFDKNEHYWVAYGLQVFDGYMAALTVACTDILMFNMISYPIGQLT
KLQHLVRNMAVYKTHFEAFPTFTKIVQRHKHVIKYVELFNQSMGTFAIFEFVQSSVQIASVLVQTSPDDLTLMSFC
FIVLFFTSMLTRLFMYYYSANEVIIQSINLGDSVWESSWYHQPHQLKQAMLMVLVRAQKPVSYKIGGFGIMSMQSI
VAILKATYTYISVILRN

>ItypOR29
MAAYPQCKNLRVAIIYSSIIGVFPWQFMFQHNHLRQTLYRWYSVFLHFWFSGFIITEYIELYLQCTADELKLDEICA
NICVVMVFTSTAVRQLVMRFNKMVNDLIQSIIDEEKHNDFLEDDKTREIEDKFIKSSDSISNWYAAPVYITLFQYVLF
PMMSKPDIIQIGNTTQALRPLIVDSWFPFDKMEYYWIVYVLQLFDLLIGALYVTYLHILMFNMYRYPVAQLKKLQHV
LRNFGRYKVEYMRQSNCNEYISALVVFRECIKKHKKIIQYVDGINECMSTYTVFDFLQSSFQIAALLVQTSPNDMTF
ISFLTVFTFITTVMIRLFVYYHSGNELIFESVNISMAIWESNWHEQSPQIKSMMLLVMRRAQKPLCYTIGGFGVMSL
QSVIAILKATYSYVSIIFRQ
